# AC-PCA: simultaneous dimension reduction and adjustment for confounding variation

**DOI:** 10.1101/040485

**Authors:** Zhixiang Lin, Can Yang, Ying Zhu, John C. Duchi, Yao Fu, Yong Wang, Bai Jiang, Mahdi Zamanighomi, Xuming Xu, Mingfeng Li, Nenad Sestan, Hongyu Zhao, Wing Hung Wong

## Abstract

Dimension reduction methods are commonly applied to high-throughput biological datasets. However, the results can be hindered by confounding factors, either biologically or technically originated. In this study, we extend Principal Component Analysis to propose AC-PCA for simultaneous dimension reduction and adjustment for confounding variation. We show that AC-PCA can adjust for a) variations across individual donors present in a human brain exon array dataset, and b) variations of different species in a model organism ENCODE RNA-Seq dataset. Our approach is able to recover the anatomical structure of neocortical regions, and to capture the shared variation among species during embryonic development. For gene selection purposes, we extend AC-PCA with sparsity constraints, and propose and implement an efficient algorithm. The methods developed in this paper can also be applied to more general settings.

## 1 Introduction

Dimension reduction methods, such as Multidimensional Scaling (MDS) and Principal Component Analysis (PCA), are commonly applied in high-throughput biological datasets to visualize data in a low dimensional space, identify dominant patterns and extract relevant features [29, 27, 8, 15, 23, 4]. MDS aims to place each sample in a lower-dimensional space such that the between-sample distances are preserved as much as possible [16]. PCA seeks the linear combinations of the original variables such that the derived variables capture maximal variance [12]. One advantage of PCA is that the principal components (PCs) are more interpretable by checking the loadings of the variables.

Confounding factors, technically or biologically originated, are commonly observed in high throughput biological experiments. Various methods have been proposed to remove the unwanted variation through regression models on known confounding factors [10], factor models and surrogate vector analysis for unobserved confounding factors [19, 6, 18, 31, 24, 28]. However, directly removing the confounding variation using these methods may introduce bias, as a result of incorrect model assumption, and it can also remove the desired biological variation. Moreover, limited work has been done in the context of dimension reduction.

To address the limitations of existing methods, we have developed AC-PCA for simultaneous dimension reduction and adjustment for confounding variation. We demonstrate the performance of AC-PCA through its application to a human brain development exon array dataset [15], a model organism ENCODE (modENCODE) RNA-Seq dataset [3, 7], and simulated data. In the human brain dataset, we found that visualization of the neocortical regions is affected by confounding factors, likely originating from the variations across individual donors (Fig.1A). As a result, a) there is no clear pattern observed among the neocortical regions and samples from the same individual donors tend to form clusters, and b) it is challenging to identify neurodevelopmental genes with interregional variation. In contrast, applying AC-PCA to the human brain dataset, we are able to recover the anatomical structure of neocortical regions and reveal temporal dynamics that existing methods are unable to capture. In the modENCODE RNA-Seq dataset, the variation across different species makes the identification of conserved developmental mechanisms challenging (Fig.3C). Our proposed method is able to capture the shared variation among species and identify genes with consistent temporal patterns in *D. melanogaster* (fly) and *C. elegans* (worm) embryonic development. We also extended AC-PCA with sparsity constraints for variable selection and better interpretation of the PCs.

**Figure 1:**
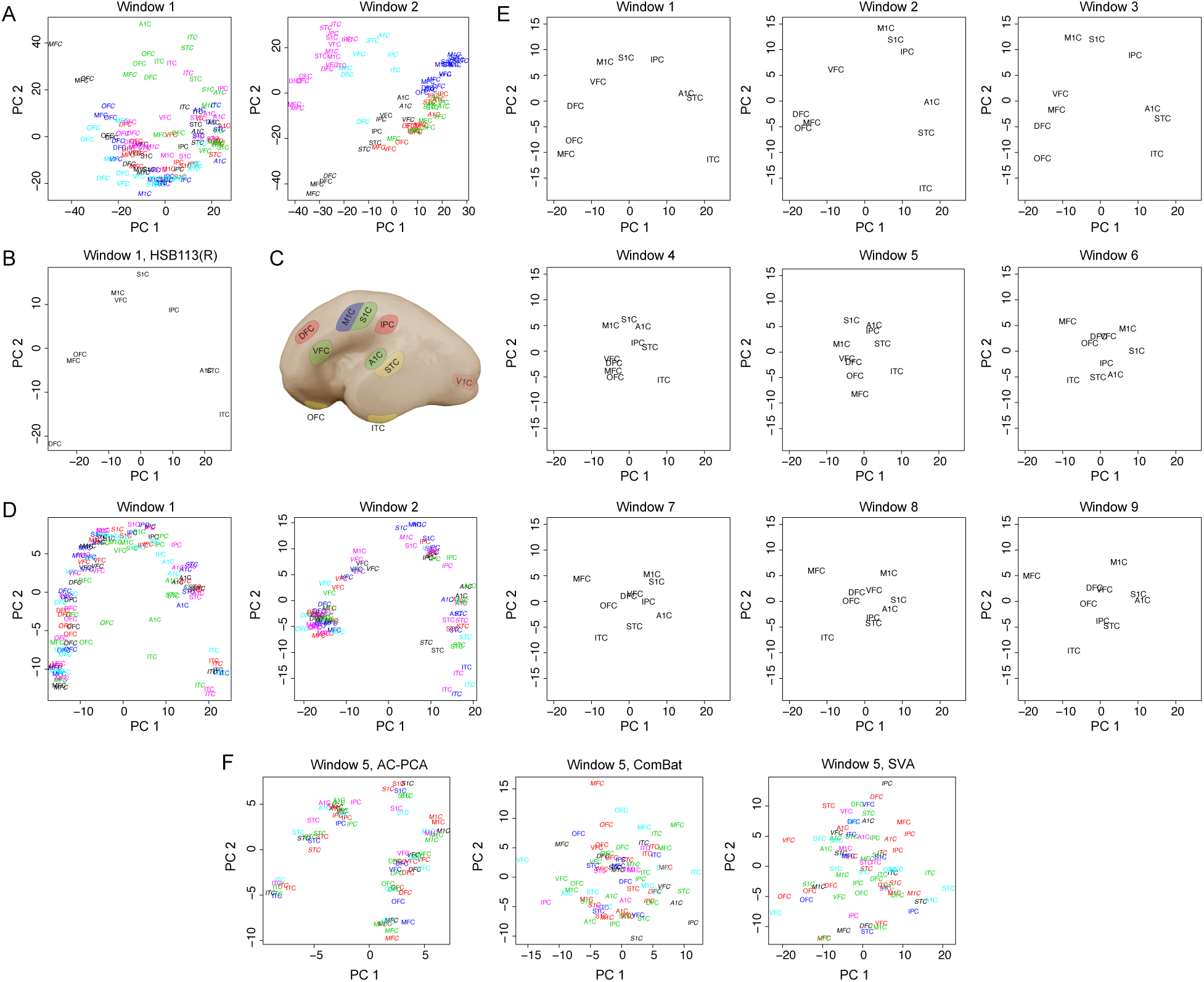
AC-PCA recovers the anatomical structure of neocortex. (*A*) PCA for time windows 1 and 2. Each color represents a donor. Samples from the right hemisphere are labeled as italic. (*B*) PCA for the right hemisphere of donor HSB113. (*C*) Representative fetal human brain, lateral surface of hemisphere[15]. MFC is not visible on the lateral surface. (*D*) AC-PCA for time windows 1 and 2. (*E*) Temporal dynamics of the principal components. Median was taken across individuals for each brain region. (*F*) Comparison between AC-PCA, ComBat[10] and SVA [19, 24], time window 5.

## 2 Results

### 2.1 Application to the human brain exon array data

The human brain exon array dataset [15] includes the transcriptomes of 16 brain regions comprising 11 areas of the neocortex, the cerebellar cortex, mediodorsal nucleus of the thalamus, striatum, amygdala and hippocampus. Because the time period system defined in [15] had varying numbers of donors across developmental epochs, beginning from period 3, we grouped samples from every 6 donors, sorted by age. While the last time window had only 5 donors, this reorganization more evenly distributed sample sizes and allowed improved comparisons across time (Table S1).

In the analysis, we used samples from 10 regions in the neocortex. V1C was excluded from the analysis as the distinct nature of this area relative to other neocortical regions tended to compress the other 10 regions into a single cluster. We conducted PCA for windows 1 and 2, as shown in Fig.1A. At first glance, neither analysis produced any clear patterns among neocortical regions. However, closer observation of these plots suggested some underlying structure: we performed PCA on just the right hemisphere of donor HSB113 and found that the gross morphological structure of the hemisphere was largely recapitulated (Fig.1B). The gross structure tends to be consistent between hemispheres and across donors within time windows when PCA is performed simultaneously, but the pattern is largely distorted, likely due to the small sample size and noisy background (Fig.S1).

In contrast, when we applied AC-PCA to see the effectiveness of our approach in adjusting confounding effects from individual donors, we were able to recover the anatomical structure of neocortex (Fig.1C and 1D). Next, we explored the temporal dynamics of the PCs (Fig.1E). The pattern is similar from windows 1 to 5, with PC1 representing the frontal to temporal gradient, which follows the contour of developing cortex [23], and PC2 representing the dorsal to ventral gradient. Starting from window 6, these two components reversed order. We also calculated the interregional variation explained by the PCs (Fig.S2). The interregional variation explained by the first two PCs decreases close to birth (window 4) and then increases in later time windows, similar to the “hourglass” pattern previously reported based on cross-species comparison and differential expression [25, 21].

We compared AC-PCA with ComBat [10] and SVA [19, 24], where PCA was implemented after removing the confounder effects. In windows 1 to 3, AC-PCA performs slightly better; In windows 4 to 9, AC-PCA outperforms ComBat and SVA (Fig.S3 and Fig.S4). The results for window 5 are shown in Fig.1F.

We then implemented AC-PCA with sparsity constrains to select genes associated with the PCs. The number of genes with non-zero loadings are shown in Fig.2A, along with the interregional variation explained in the regular PCs. Interestingly, the trends tend to be consistent: when the regular PC explains more variation, more genes are selected in the corresponding sparse PC. To produce more stringent and comparable gene lists, we chose the sparsity parameter such that 200 genes are selected in each window. The overlap of gene lists across windows is moderate (Fig.S5) and, as expected, the overlap with the first window decreases over time. The overlap between adjacent windows tends to be larger in later time windows, indicating that interregional differences become stable. In windows 1 and 3, genes with the largest loadings demonstrate interesting spatial patterns (Fig.2B). For PC1, the top genes follow the frontal to temporal gradient; while for PC2, they tend to follow the dorsal to ventral gradient. A brief overview of the functions of these genes is shown in Table S2.

**Figure 2:**
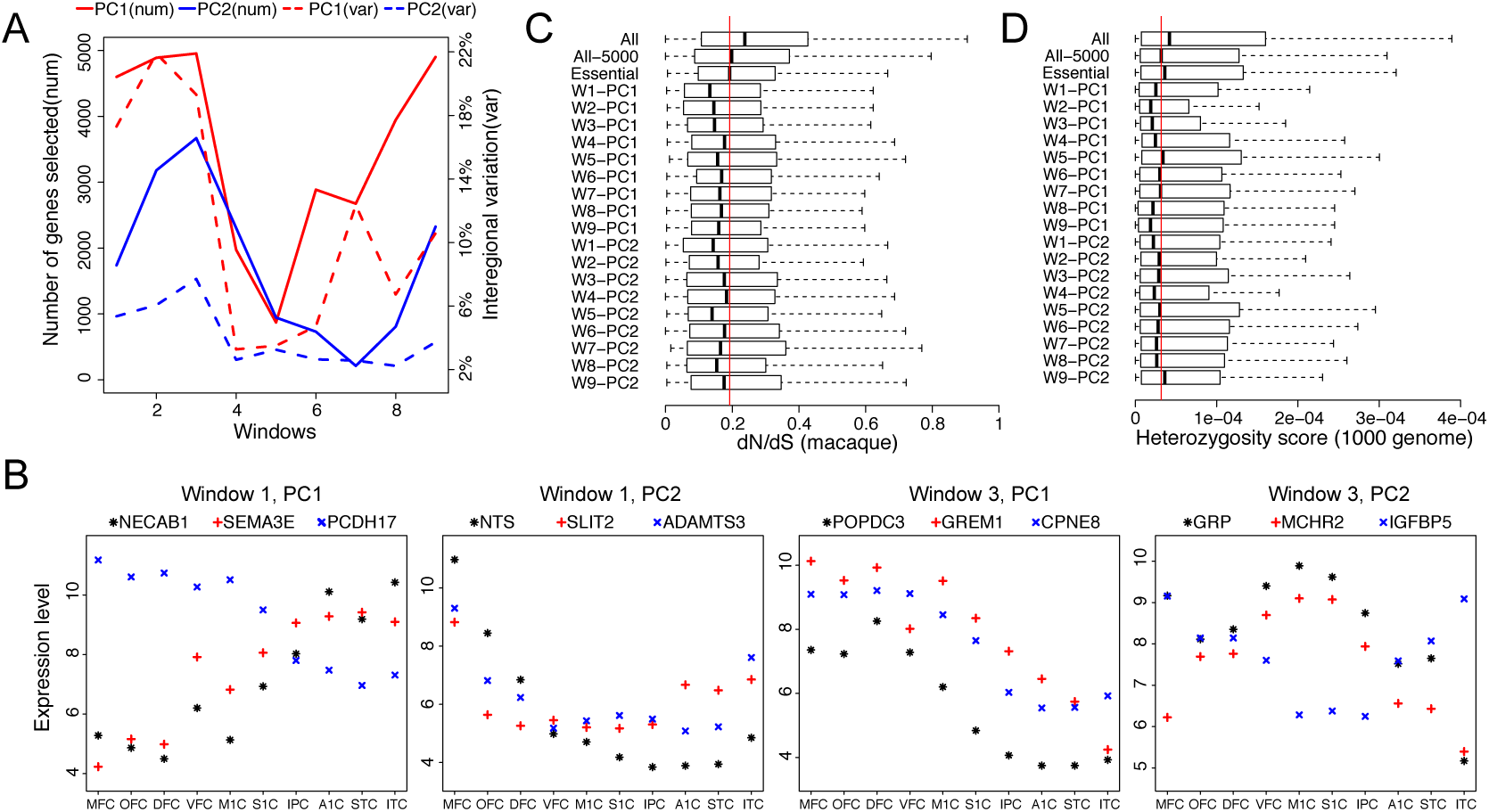
Gene selection by AC-PCA with sparse loadings. (*A*) Number of genes selected in the sparse PCs and the interregional variation explained by the regular PCs. Interregional variation is calculated to be the sum over the variance across regions among the individuals. (*B*) Expression levels for the genes with top loadings in windows 1 and 3. The top 3 genes are shown and each point represents the median over the individuals. (*C*) and (*D*) Conservation (dN/dS) and heterozygosity scores for the genes in PC1 and PC2. “All” represents the total 17, 568 genes in the exon array experiment. “All-5000” represents the 5,000 genes used in the analysis. “Essential” represents a list of 337 genes that are functionally conserved and essential, obtained from the Database of Essential Genes (DEG) version 5.0 [32]. “Window” is abbreviated as “W”.

Finally, we demonstrate the functional conservation of the 200 genes selected in PC1 and PC2. These genes tend to have low dN/dS scores for human vs. macaque comparison, even lower than the complete list of all essential genes (Fig.2C). In the human vs. mouse comparison, we observed a similar trend (Fig.S6). Parallel to the cross-species conservation, we also observed that these genes tend to have low heterozygosity scores, a measure of functional conservation in human (Fig.2D).

### 2.2 Application to the modENCODE RNA-Seq data

The modENCODE project generates the transcriptional landscapes for model organisms during development[3, 7]. In the analysis, we used the time-course RNA-Seq data for fly and worm embryonic development.

We first conducted PCA on fly and worm separately, as shown in Fig.3A. Although the temporal patterns share some similarity, the projections for fly and worm are different. The genes with top loadings in fly have different temporal dynamics in worm, especially for PC2 (Fig.3B).

**Figure 3:**
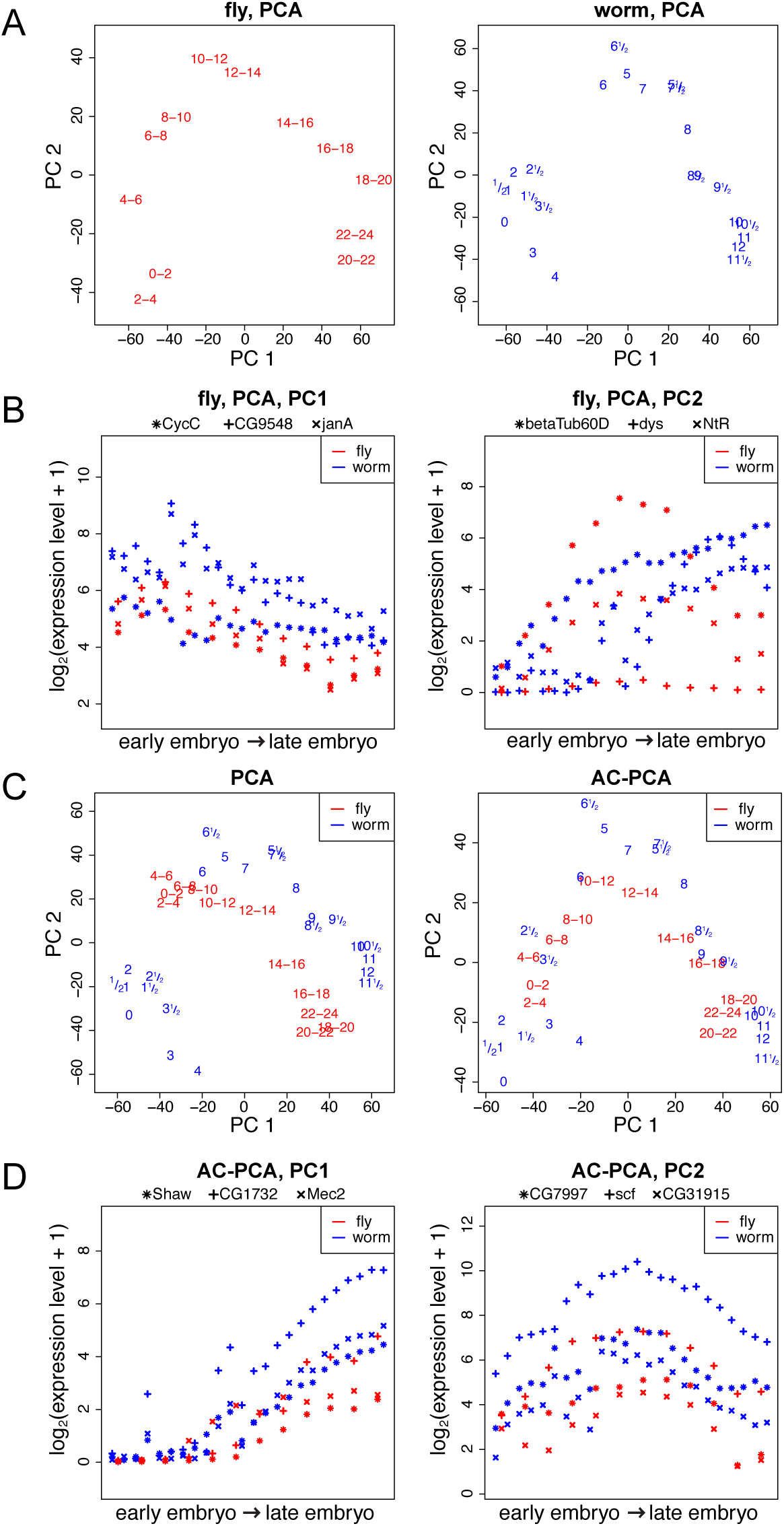
AC-PCA captures shared variation in fly and worm modENCODE RNA-Seq data, embryonic stage. (*A*) PCA on fly and worm separately. Each data point represents a time window (fly) or a time point (worm), in the unit of hours. (*B*) Expression levels of the top 3 genes in fly PCA. (*C*) PCA and AC-PCA on fly and worm jointly. (*D*) Expression levels of the top 3 genes in AC-PCA.

We also conducted PCA on fly and worm jointly, which reveals the variations of different species (Fig.3C). We demonstrate the performance of AC-PCA in capturing the shared variation among fly and worm (Fig.3C). The selected genes tend to have consistent and smooth temporal patterns in both species (Fig.3D). PCA on fly and worm jointly cannot capture the direction of PC2 in AC-PCA, which the gene expression levels peak in middle embryonic stage.

### 2.3 Simulations

We tested the performance of AC-PCA on simulated datasets with various settings. The number of variables *p* = 400. More details on the simulation are provided in the Materials and Methods Section. We first considered simulations with individual variation:

Setting 1: individual variation is represented by a global trend for all the variables.
Setting 2: for some variables, the variation is shared among individuals, and it is not shared for the other variables. We also considered other confounding structure:
Setting 3: the data is confounded with two experimental “batches”, each contributing globally to the data.
Setting 4: the data is confounded with a continuous confounding factor (e.g. age), contributing globally to the data. The results are presented in Fig.4. In all simulations, AC-PCA is able to recover the true pattern. Finally, we tested AC-PCA for variable selection when the true loading is sparse:
Setting 5: similar to setting 2, except that the true loading is set to be sparse, and for simplicity, we assumed that the latent factor is of rank 1.Results for simulation setting 5 are shown in Table 1, where *α* indicates the magnitude of confounding variation, and *σ* indicates the noise level. Larger noise leads to lower sensitivity, larger standard error for the estimated non-zeroes, but does not affect the mean much. Smaller confounding variation leads to overestimate of the non-zeroes, but does not affect the sensitivity much.

**Figure 4:**
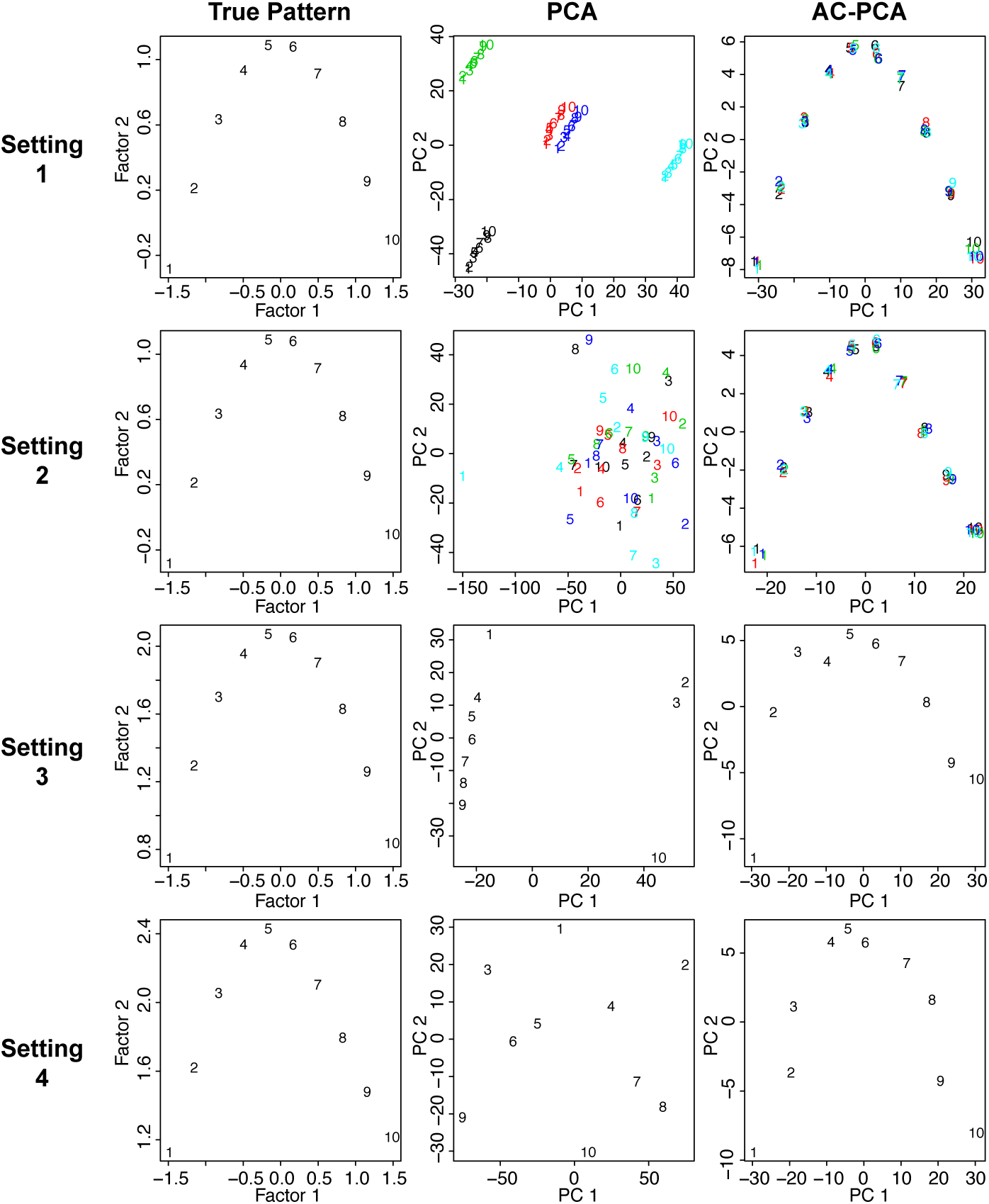
Visualization of simulated data with confounding variation.

**Table 1:** Sparsity estimation and sensitivity

|  |  | Estimated non-zeros |  | Sensitivity<sup>‡</sup> |  |
| --- | --- | --- | --- | --- | --- |
| | | $\sigma = 0.2$ | $\sigma = 0.5$ | $\sigma = 0.2$ | $\sigma = 0.5$ |
| 100* | $\alpha = 1.5$ | 113.7(34.2) | 120.6(42.5) | 0.85(0.06) | 0.80(0.07) |
| | $\alpha = 2.0$ | 88.7(20.9) | 91.6(31.1) | 0.83(0.08) | 0.79(0.08) |
| | $\alpha = 2.5$ | 81.4(20.5) | 79.4(24.4) | 0.83(0.08) | 0.80(0.05) |
| 40* | $\alpha = 2.0$ | 37.4(14.7) | 38.9(15.6) | 0.74(0.11) | 0.62(0.10) |
| | $\alpha = 2.0^\dagger$ | 46.7(16.5) | 54.6(19.0) | 0.77(0.10) | 0.69(0.11) |

\* The true number of non-zeros; <sup>†</sup> The true variation was scaled with 2.5 to let its magnitude match with that in the less sparse setting; <sup>‡</sup> To calculate sensitivity, the sparsity parameter $c_2$ was chosen such that the estimated non-zero entries equals the true number.

## 3 Discussion

Dimension reduction methods are commonly applied to visualize high-dimensional data in a lower dimensional space and to identify dominant patterns in the data. Confounding variation may affect the performance of these methods, and hence the visualization and interpretation of the results (Fig. 1A and 3C).

In this study, we have proposed AC-PCA for simultaneous dimension reduction and adjustment for confounding variation. One feature of AC-PCA is its simplicity. Instead of specifying analytical forms for the confounding variation, we extended the objective function of regular PCA with penalty on the dependence between the PCs and the confounding factors. AC-PCA is designed to capture the desired biological variation, even when the confounding factors are unobserved, as long as the labels for the primary variable of interest are known.

The application of AC-PCA is not limited to transcriptome datasets. Dimension reduction methods have been applied to other types of genomics data for various purposes, such as feature extraction for methylation prediction [5], classifying yeast mutants using metabolic footprinting [1], classifying immune cells using DNA methylome [17], and others. AC-PCA is applicable to these datasets to capture the desired variation, adjust for potential confounders, and select the relevant features. AC-PCA can serve as an exploratory tool and be combined with other methods. For example, the extracted features can be implemented in regression models. The R package and Matlab source code with user’s guide and application examples is available on https://github.com/linzx06/AC-PCA.

## 4 Methods

### 4.1 AC-PCA adjusting for variations of individual donors

Let *X*_*i*_ represent the *b* × *p* matrix for the gene expression levels of individual *i*, where *b* is the number of brain regions and *p* is the number of genes. By stacking the rows of *X*_1_,…,*X*_*n*_, *X* represents the (*n* × *b*) × *p* matrix for the gene expression levels of *n* individuals. We propose the following objective function to adjust for individual variation:

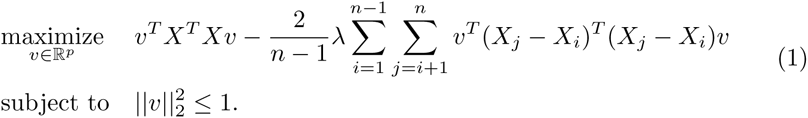

In (1), the term *υ*^*T*^*X*^*T*^*X*_*υ*_ is the objective function for standard PCA, and the regularization term 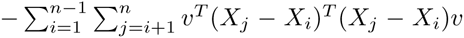 encourages the coordinates of the brain regions across individuals to be similar. The factor 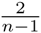 makes the regularization term in formulation (1) scale linearly with the number of individuals. The tuning parameter λ > 0 controls the strength of regularization. When λ = + ∞, we are forcing the coordinates of the same brain region across individuals to be the same after projection. Only the labels for brain regions (i.e. labels for the primary variables) are required when implementing formulation (1). We can apply it even if the individual labels of donors (i.e. the confounding variables) are unknown. The connection of formulation (1) with Canonical Correlation Analysis (CCA) is shown in SI Materials and Methods.

### 4.2 AC-PCA capturing shared variations among species

For the fly data, samples were taken in 12 time windows during embryonic development: 0-2, 2-4,…, 22-24 hours; For the worm data, samples were taken every 30 minutes during embryonic development: 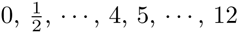 hours, where the sample from 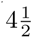 hours is missing, resulting in 24 samples. Let *X*^*f*^ and *X*^*w*^ represent the data matrices for fly and worm, correspondingly. Let *X*_*f*_ and *X* represent the data matrix for both species, by stacking the rows in *X*^*f*^ and *X*^*w*^. We propose the following objective function to adjust for the variation of species:

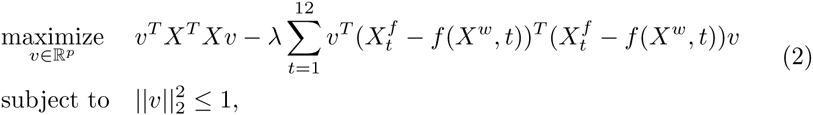

where

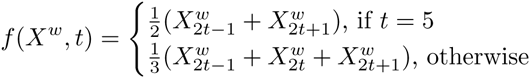

Without prior knowledge on the alignment of fly and worm developmental stages, it is reasonable to shrink the projection of 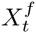 towards the mean of 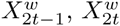 and 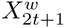 after projection. [20] aligned fly and worm development based on stage-associated genes using the same dataset. We did not implement AC-PCA with their alignment results, to keep the analysis unsupervised.

### 4.3 AC-PCA in a general form

Let *X* be the *N* × *p* data matrix and *Y* be the *N* × *l* matrix for *l* confounding factors. Denote *y*_*i*_ the *i*th row in *Y*. We propose the following objective function to adjust for more general confounding variation:

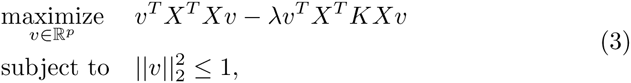

where *K* is the *N* × *N* kernel matrix, and *K*_*ij*_ = *k*(*y*_*i*_, *y*_*j*_). It can be shown that *υ*^*T*^*X*^*T*^*KX*_*υ*_ is the same as the empirical Hilbert-Schmidt independence criterion[9, 2] for *X*_*υ*_ and *Y*, where linear kernel is applied on *X*_*υ*_ (SI Materials and Methods). In the objective function, we are penalizing the dependence between projected data *X*_*v*_ and the confounding factors *Y*. Formulations (1) and (2) are special cases of (3). In formulation (1), linear kernel (i.e. *YY*^*T*^) is applied on *Y*, and *Y* has the following structure: in each column of *Y*, there are only two non-zero entries, 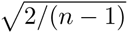 and 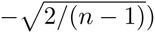, corresponding to a pair of samples from the same brain region but different individuals. Denote *Z* = *X*^*T*^*X* — λ*X*^*T*^*KX*. Problem (3) can be rewritten as:

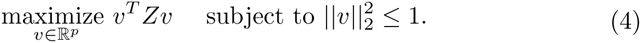

Therefore it can be solved directly by implementing eigendecomposition on *Z*.

### 4.4 AC-PCA with sparse loading

In PCA, the loadings for the variables are typically nonzero. In high dimensional settings, sparsity constraints have been proposed in PCA for better interpretation of the PCs [13, 33, 30, 26] and better statistical properties, such as asymptotic consistency [11, 14, 22].

Denote *H* = *X*^*T*^*KX*. It can be shown that solving (3) is equivalent to solving:

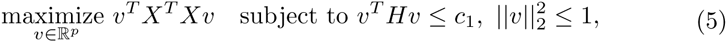

where *c*_1_ is a constant depending on λ. A sparse solution for *υ* can be achieved by adding *ℓ*_1_ constraint:

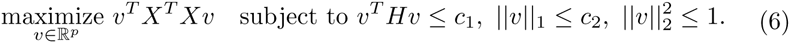

Following [30], this is equivalent to:

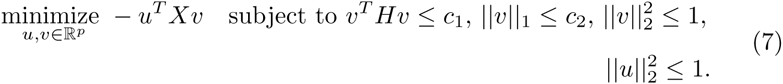

Problem (7) is biconvex in *u* and *υ*, and it can be solved by iteratively updating *u* and *υ*. At the *k*th iteration, the update for *u* is simply 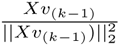. To update *υ*, we need to solve:

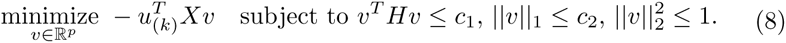

Because of the quadratic constraint on *υ*, it is hard to solve (8) directly. We propose to use the bisection method to solve the following feasibility problem iteratively:

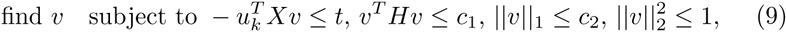

where *t* is an upper-bound for 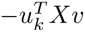 and is updated in each iteration. Problem (9) can be solved by alternating projection. Details for the algorithm are included in the SI Materials and Methods.

### 4.5 Multiple principal components

In (3), obtaining multiple principal components is straightforward, as they are just the eigenvectors of *Z*; for the sparse solution, (7) can only obtain the first sparse principal component. To obtain the other sparse principal components, we can update *X* sequentially with 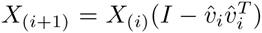 for *i* = 1, …, and implement (7) on *X*_(*i*+1)_, where *X*_(1)_ = *X* and *υ̂*_*i*_ is the *ith* principal component.

### 4.6 Tuning λ

Let *X* denote the *N* × *p* data matrix. *l* = *υ*^*T*^*X*^*T*^*KXυ* can be treated as a loss function to be minimized. We do 10-fold cross-validation to tune λ:

1. From *X*, we construct 10 data matrices *X*_1_,…,*X*_10_, each of which is missing a non-overlapping one-tenth of the rows in *X*
2. For each *X*_*i*_, *i* =1,…, 10, implement (3) and obtain *υ*_*i*_
3. Calculate *l*_cυ_. In *X*_*υ*_, we use *υ*_*i*_ and the missing rows that are left out in *X*_*i*_, for *i* = 1,…, 10
4. When λ increases from 0, *l*_*cυ*_ usually decreases sharply and then either increases or becomes flat. In practice, we choose λ to be the “elbow” point of *l*_*cυ*_.

In simulations and real data analysis, the overall patterns recovered by AC-PCA are robust to the choice of λ. In the human brain dataset, every region tends to form a smaller cluster, variation of the principal components shrinks, but the overall pattern remains similar, when λ is larger than the best tuning value (Fig.S7). For the comparison of interregional variations over the time windows, we fixed λ to the same value.

### 4.7 Tuning *c*_1_ and *c*_2_

Because *c*_1_ and *c*_2_ capture different aspects of the data, we propose a two-step approach: first tune *c*_1_, and then tune *c*_2_ with *c*_1_ fixed. This also greatly reduces the computational cost since tuning *c*_2_ can be slow. To tune *c*_1_:

1. We follow the previous procedure to tune λ
2. The best λ is used to calculate *v*,as in (3)
3. Let *c*_1_ = *υ*^*T*^*X*^*T*^*KX*_*υ*_

To tune *c*_2_, we follow Algorithm 5 in [30], which is based on matrix completion:

1. From *X*, we construct 10 data matrices *X*_1_,…,*X*_10_, each of which is missing a non-overlapping one-tenth of the elements of *X*
2. For *X*_1_,…, *X*_10_, fit (7) and obtain *X*̂_*i*_ = *duυ*^*T*^, the resulting estimate of *X*_*i*_ and *d* = *u*^*T*^*X*_*i*_*υ*
3. Calculate the mean squared errors of *X*̂_*i*_, for *i* = 1,…, 10, using only the missing entries
4. Choose *c*_2_ that minimizes the sum of mean squared errors

### 4.8 Simulations

Setting 1: we considered *n* = 5, *b* =10 and *p* = 400. For the ith individual, the *b* × *p* matrix *X*_*i*_ = *Wh* + *αBs*_*i*_ + *ϵ*_*i*_. *Wh* represents the shared variation across individuals. *αBs*_*i*_ corresponds to individual variation and *ϵ*_*i*_ is noise. *W* = (*w*_1_ *w*_2_) is a *b* × 2 matrix, representing the latent structure of the shared variation. For visualization purpose, we assumed that it is smooth and has rank 2. Let *μ* = (1,…, b)′ and *w*_1_ is the normalized *μ*, with mean 0 and variance 1. *w*_2_ ~ *N*(0, 0.25 · Σ), where 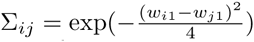. *h* is a 2 × *p* matrix and the rows in *h* are generated from *N*(0, *I*_*p*_), where *I*_*p*_ is the *p* × *p* identity matrix. *B* is a *b* × 1 matrix with all 1s. *s*_*i*_ is generated from *N*(0,*I*_*p*_). *α* is a scalar indicating the strength of confounding variation, we set *α* = 2.5. The rows in *ϵ*\* are generated from *N*(0,0.25 · *I*_*p*_).

Setting 2: we considered *n* = 5, *b* =10 and *p* = 400. For the *i*th individual, let 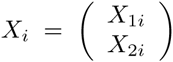, where *X*_1*i*_ represents the data matrix for the first 200 variables and *X*_2*i*_ represents that of the other 200 variables. *X*_1*i*_ = *Wh* + *ϵ*_1*i*_ and *X*_2*i*_ = *αW*_*i*_*h*_*i*_ + *ϵ*_2*i*_. *W*, *h*, *h*_*i*_, *ϵ*_1*i*_ and *ϵ*_2*i*_ are generated similarly as that in setting 1. *α* = 2.5. The first column in *W*_*i*_ is generated from *N*(0, *I*_*b*_), and the second column is generated from *N*(0, 0.25 · *I*_*b*_), where *I*_*b*_ is the *b* × *b* identity matrix.

Settings 1 and 2 represent data with individual variation and we implemented formulation 1.

Setting 3: *N* =10 and *p* = 400. The *N* × *p* matrix *X* = *Wh* + *αBs* + *ϵ*. *W* and *h* are the same as that in setting 1. We set *α* = 2.5. *B* = (*b*_1_ *b*_2_) is a *N* × 2 matrix: the entries in *b*_1_ have 0.3 probability of being 0, otherwise the entries are set to 1; *b*_2_ equals 1 – *b*_1_. *s* is a 2 × *p* matrix, where the rows are generated from *N*(0, *I*_*p*_). *ϵ* is an *N* × *p* matrix, where the rows are generated from *N*(0,0.25 · *I*_*p*_).

Setting 4: *N* =10 and *p* = 400. The *N* × *p* matrix *X* = *Wh* + *αw̃*_1_*s* + *ϵ*. *W*, *h* are the same as that in setting 1. We set *α* = 2.5.*w̃*_1_ is a permutation of *w*_1_, representing a continuous confounder. *s* is generated from *N*(0,*I*_*p*_). *ϵ* is an *N* × *p* matrix, where the rows are generated from *N*(0,0.25 · *I*_*p*_).

When implementing formulation (3), *Y* were set to *B* and *w̃*_*1*_ in settings 3 and 4,correspondingly.

Setting 5: for the ith individual, let 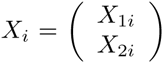, where *X*_1*i*_ represents thedata matrix for the first 200 variables and *X*_2*i*_ represents that of the other 200 variables. *X*_1*i*_ = *wh* + *ϵ*_1*i*_ and *X*_2*i*_ = *αW*_*i*_*h*_*i*_ + *ϵ*_2*i*_. *w* is the same as *w*_1_ in setting 1. Some entries in *h* are set to 0 to reflect sparsity, the other entries are generated from *N*(0,*I*). *W*_*i*_ and *h*_*i*_ are the same as that in setting 2. The rows in *ϵ*_1*i*_ and *ϵ*_2*i*_ are generated from *N*(0, *σ*^2^ · *I*), where *σ* is a scalar indicating the noise level.

### 4.9 Data preprocessing

The human brain exon array dataset was downloaded from the Gene Expression Omnibus (GEO) database under the accession number GSE25219. The dataset was generated from 1,340 tissue samples collected from 57 developing and adult post-mortem brain[15]. Details for the quality control (QC) of the dataset were described as in [15]. After the QC procedures, noise reduction was accomplished by removing genes that were not expressed [21], leaving 13,720 genes in the dataset. We next selected the top 5,000 genes sorted by coefficient of variation. The modENCODE RNA-Seq dataset was downloaded from https://www.encodeproject.org/comparative/transcriptome/. We used samples from the embryonic stage of worm and fly. The coding genes with orthologs in both worm and fly were used. For the orthologs in fly that map to multiple orthologs in worm, we took median to get a one to one match, resulting in 4S31 ortholog paris. To gain robustness, we used the rank across samples within the same species. The rank matrix was then scaled to have unit variance.

### 4.10 ComBat and SVA

ComBat and SVA were implemented with the sva package from Bioconductor. For ComBat, the input for batch covariate was the individual label, and the input for model matrix was the brain region label. For SVA, the input for model matrix was the brain region label; Functions *sva* and *f sva* were used to remove the confounding variation.

## 5 Acknowledgement

Z.L. and H.Z. were partially supported by the National Science Foundation grant DMS-1106738 and the National Institutes of Health grants R01 GM59507 and P01 CA154295; Z.L., Y.W., B.J., M.Z. and W.H.W. were partially supported by the National Institutes of Health grants R01 HG007S34 and R01 GM109S36; We thank Matthew W. State for the partial financial support of Z.L.; C.Y. was supported by grant No.61501389 from National Science Funding of China, grant No.22302815 from the Hong Kong Research Grant Council, and grant FRG2/14-15/069 from Hong Kong Baptist University the Hong Kong RGC grant HKBU; Y.Z., X.X., M.L. and N.S. were supported by the National Institutes of Health grants P50 MH106934 and U01 MH103339. All computations were performed on the Yale University Biomedical High Performance Computing Center.

## Supporting Information (SI)

Zhixiang Lin, Can Yang, Ying Zhu, John C. Duchi, Yao Fu, Yong Wang, Bai Jiang, Mahdi Zamanighomi, Xuming Xu, Mingfeng Li, Nenad Sestan, Hongyu Zhao* & Wing Hung Wong*

April 18, 2016

### SI Materials and Methods

#### Conservation and heterozygosity scores

The dN/dS score for cross-species conservation was calculated using Ensembl BioMart [11]. The heterozygosity score was calculated using 1,000 Genomes phase 1 version 3 [9]. Let *f*_1_,…*f*_*p*_ denote the allele frequencies for the *p* non-synonymous coding SNPs in a gene, and let *l* denote the maximum transcript length over all isoforms of that gene. The heterozygosity score was calculated as: 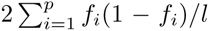. For a gene with low heterozygosity score, the non-synonymous variants in that gene tend to be rare, which indicates the functional importance of that gene.

#### Connection with Canonical Correlation Analysis (CCA)

Without loss of generality, let *n* = 2, then the objective function becomes:

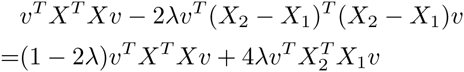

In Canonical Correlation Analysis (CCA), there are two datasets *X* and *Y*, and the goal is to maximize the correlation between *X* and *Y* after projections. The objective function in CCA is *a*^*T*^*X*^*T*^*Yb*, where *a* and *b* are two column vectors. Note that *a*^*T*^*X*^*T*^*Yb* and 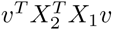 have similar forms: in 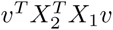 the projection vectors *a* and *b* are the same as *v*. If we treat the data from the two individuals as separate datasets, the objective function mimics a balance between regular PCA and CCA. When λ > 0.5, the weight on PCA is negative.

#### Hilbert-Schmidt independence criterion

The linear kernel of *X*_*υ*_ is *L* = *X*_*υυ*_^*T*^X^*T*^. Let *K* be the kernel of *Y*. Let *H* = *I* – *N*^‒1^ *ee*^*T*^, where *e* is a column vector with all 1s. Then *H*^*T*^*X* = *HX* = *X*, as *X* is centered. The empirical Hilbert-Schmidt independence criterion for *Xυ* and *Y* is:

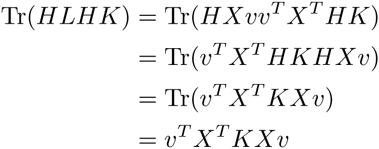

#### Details for the bisection method

The bisection method solves the following feasibility problem iteratively:

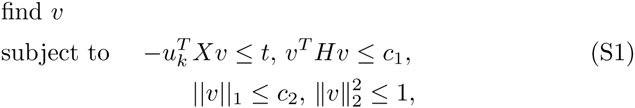

where *t* is an upper-bound for 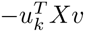 and is updated in each iteration. Algorithm 1: Bisection method for updating *v*

1. Initialize *t*_*up*_ = 0 and *t*_*low*_ = —||*X*^*T*^*u*_*k*_||_2_
2. Iterate until convergence:

(a) *t^*^* = ^(*t*^*up* + *t*_*low*_)/2
(b) *t*_*up*_ ← *t*^*^ if (*S* 1) is feasible for *t*^*^
(c) *t*_*low*_ ← *t*^*^ if (S1) is not feasible for *t*_*_
3. Let *t* = *t*_*up*_ and find *υ* by solving (S1)

The feasibility problem (S1) can be solved by alternating projection on the convex sets:

#### Projection onto hyperplane

Finding the projection of *v*_*0*_ onto the hyperplane:

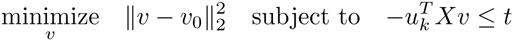

The solution is:

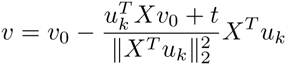

#### Projection onto *l*_2_ ball

First do decomposition *K* = ΔΔ^*T*^, and let *M* = Δ^*T*^*X*. We have *M*^*T*^*M* = *X*^*T*^ΔΔ^*T*^*X* = *X*^*T*^*KX*. We need to solve the following optimization problem:

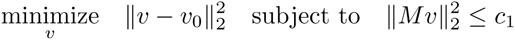

It is equivalent to the following Lagrangian problem:

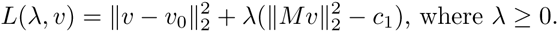

For any fixed λ, *L*(λ, *v*) is a convex differentiable function of *υ*. By taking derivative, it can be shown that *υ*_*_ ≡ *υ*_*_(λ) = (*I*_*p*_ + λ*M*^*T*^ *M*)^‒1^ *υ*_0_ minimizes the Lagrangian. Then we have

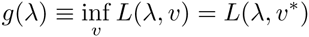

By singular value decomposition, *M* = *U*Σ*V*^*T*^, where *U*^*T*^*U* = *UU*^*T*^ = *I*_*N*_ and *V*^*T*^ *V* = *VV*^*T*^ = *I*_*p*_. Then we have

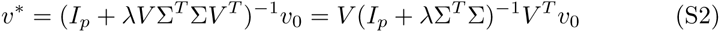

and

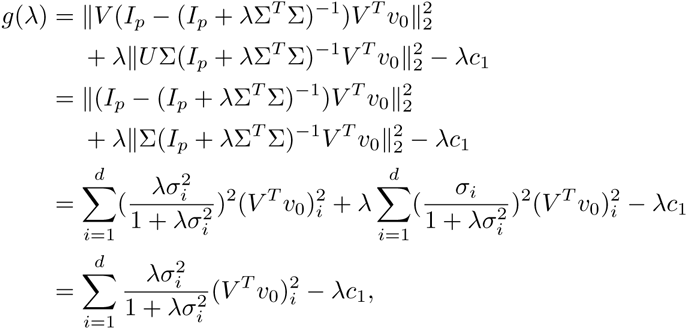

where *d*(<= *N*) is the number of non-zero singular values of *M*,*iα*_*i*_ is the *i*th non-zero singular value, and (·)*i* represents the *i*th element of a vector. When λ≥0, g(λ) is concave and the optimal value λ_*\**_ can be found by Newton-Raphson’s Method. With λ = λ_*_, the projection *υ* can be calculated with (S2), where the inversion part is a diagonal matrix. It can be shown that only the first *d* columns of *V* affect the projection and typically we have *d* ≪ *p*.

#### Projection onto *l_1_* ball

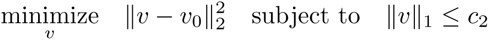

This can be solved efficiently by the algorithm presented in [12].

**Projection onto *ℓ*_2_ ball-2**

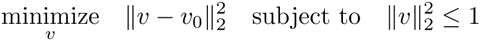

The solution is:

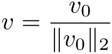

### SI Figures

**Figure 1:**
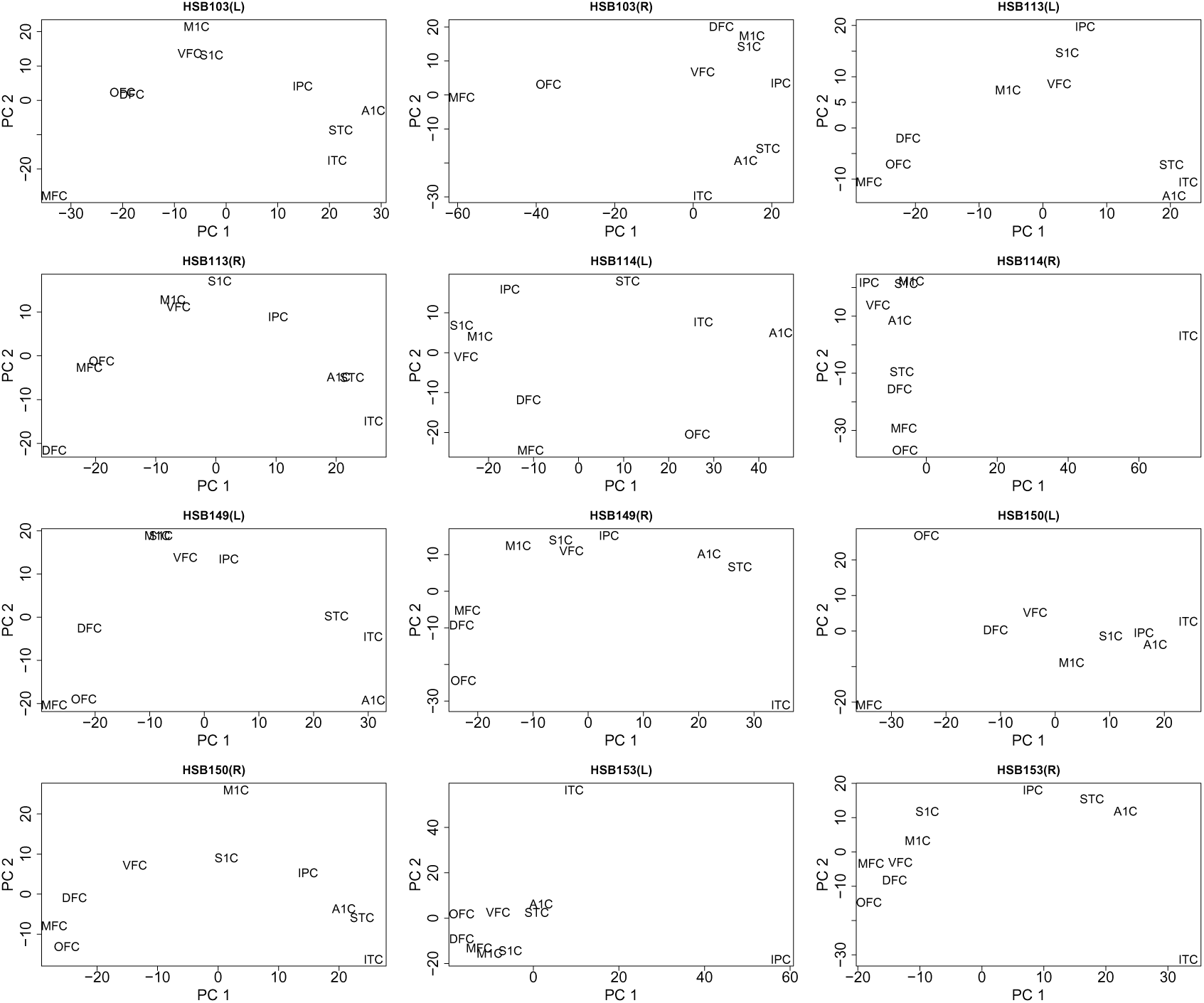
PCA for all the individuals in window 1. “R” represents the right hemisphere, and “L” represents the left hemisphere.

**Figure 2:**
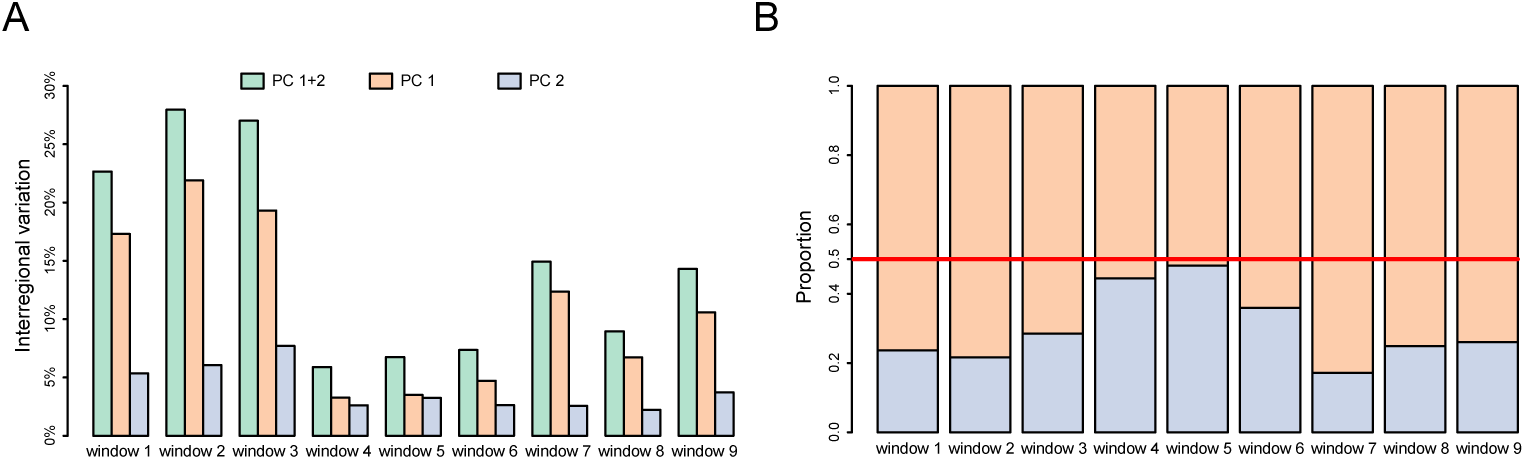
Temporal dynamics of the interregional variation. For each individual, the variance across regions was first calculated; Interregional variation equals the sum of variance over individuals. (*A*) Interregional variation explained by the first two PCs in AC-PCA. (*B*) Proportion of variation explained by the first two PCs in AC-PCA.

**Figure 3:**
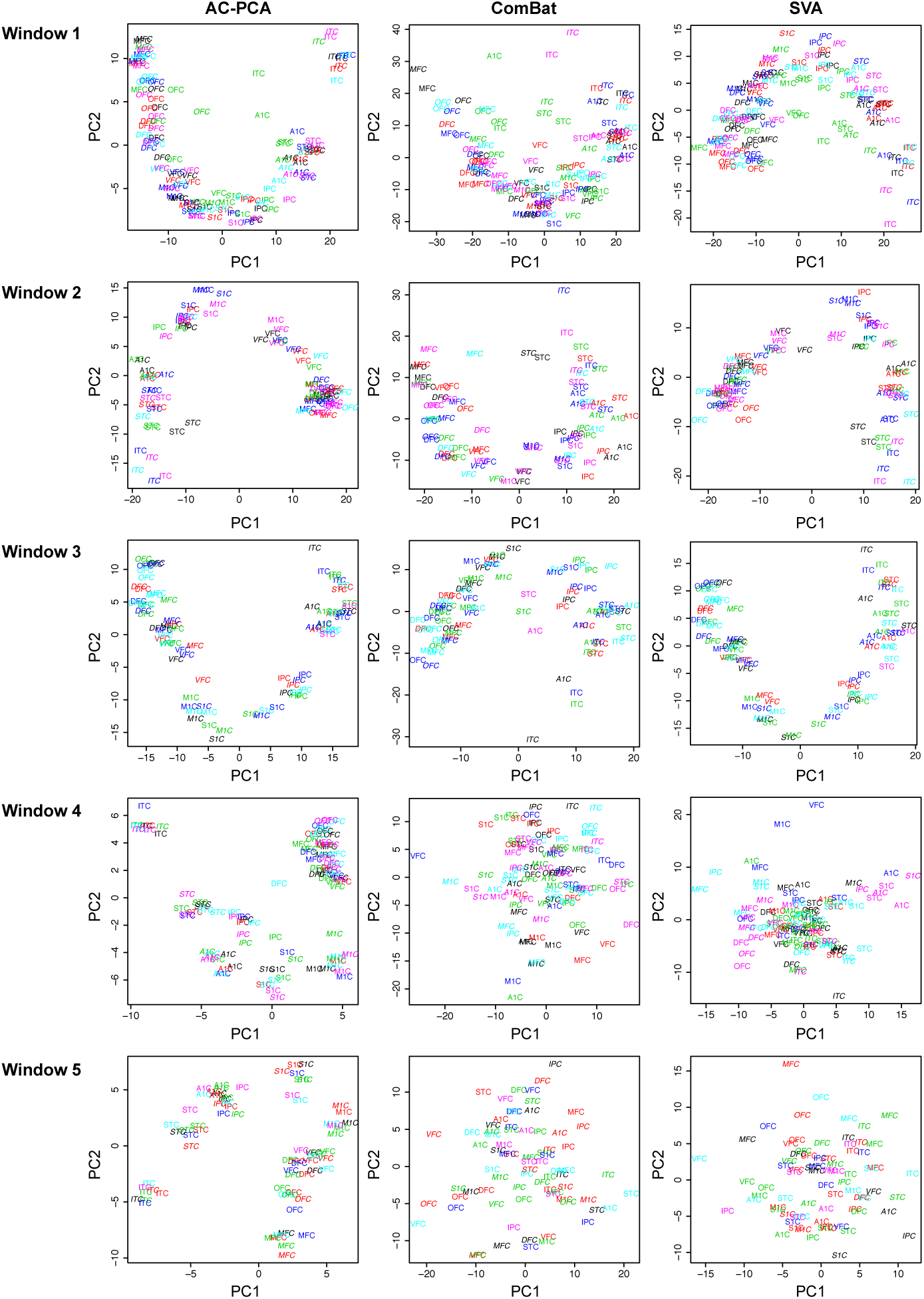
Comparison of AC-PCA, ComBat and SVA, windows l to 5. For ComBat [l8] and SVA [20, 22], PCA was implemented after removing the confounder effects.

**Figure 4:**
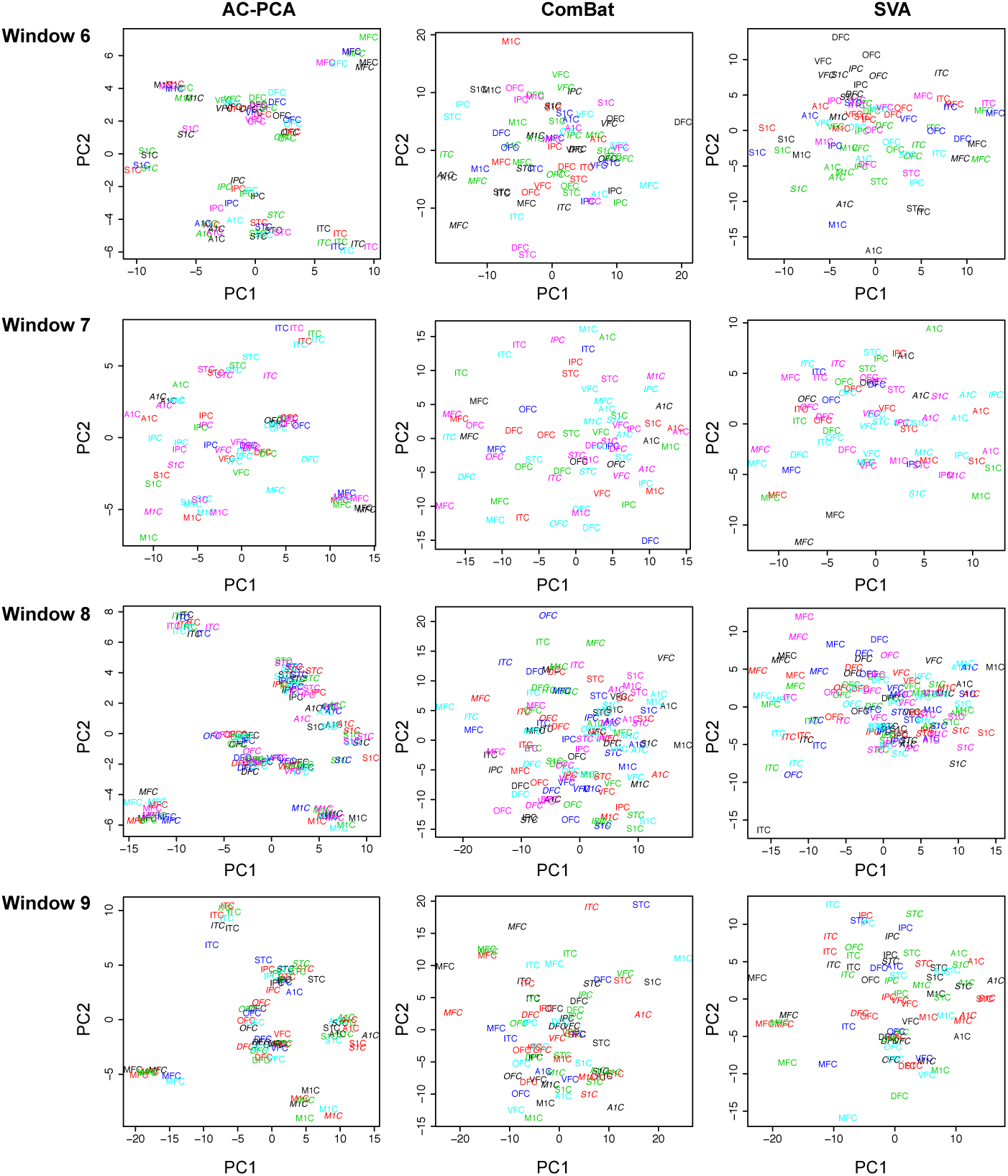
Comparison of AC-PCA, ComBat and SVA, windows 6 to 9. For ComBat [lSj and SVA [20, 22], PCA was implemented after removing the confounder effects.

**Figure 5:**
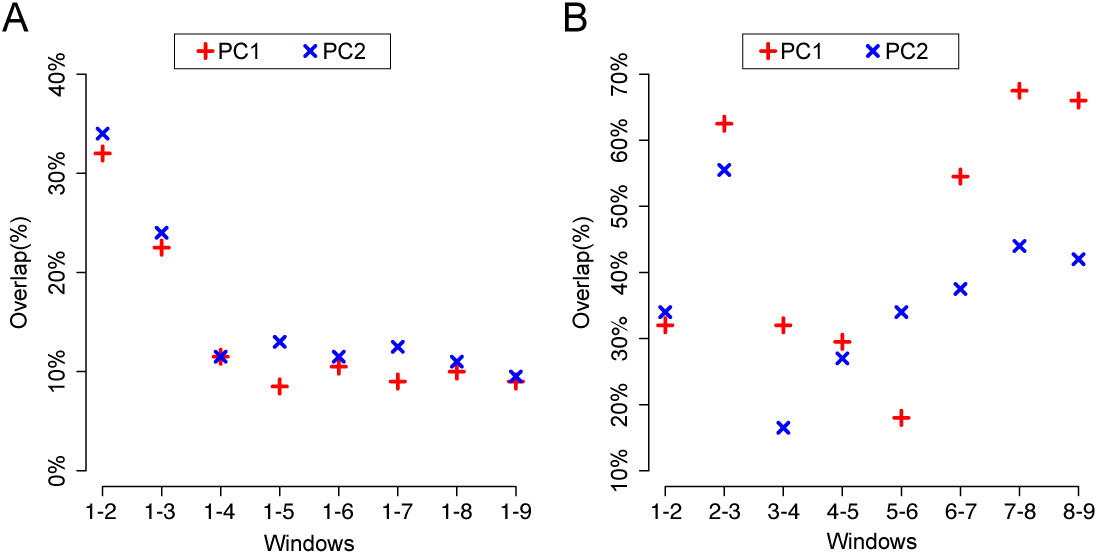
Overlap of the top 200 genes in PC1 and PC2. (*A*) Overlap with window 1. (*B*) Overlap between adjacent windows

**Figure 6:**
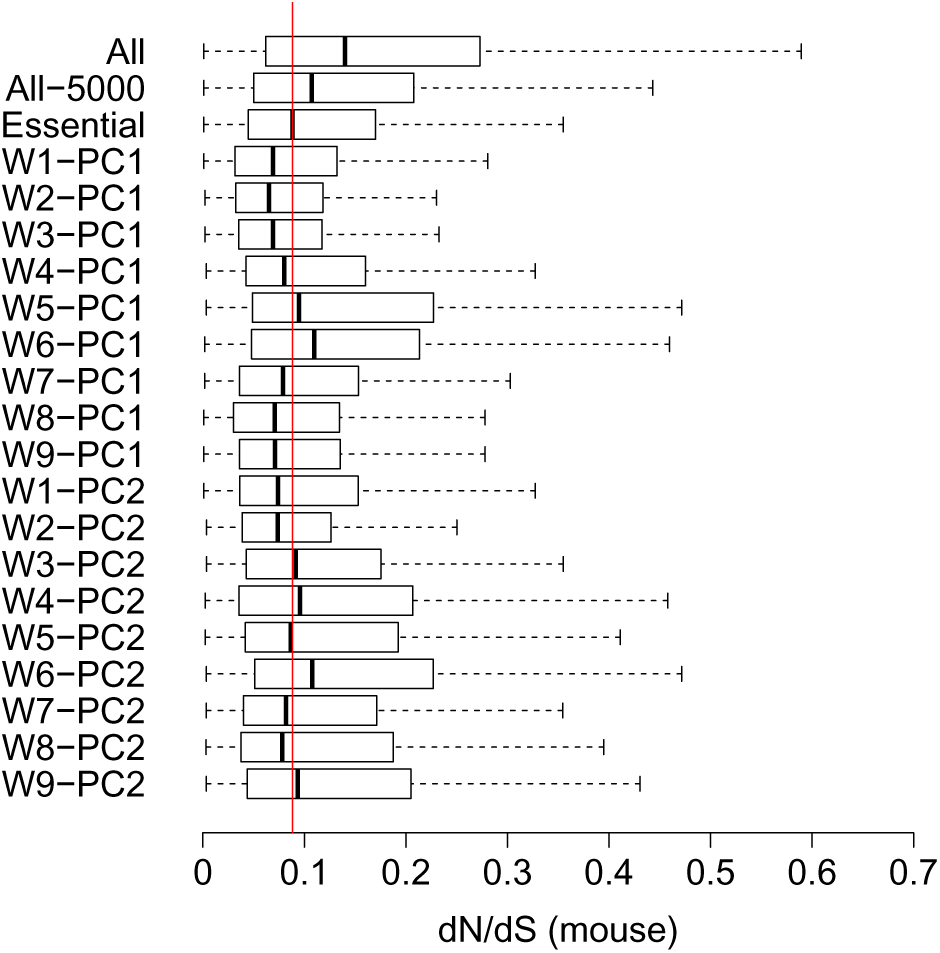
Conservation scores for the human vs. mouse comparison

**Figure 7:**
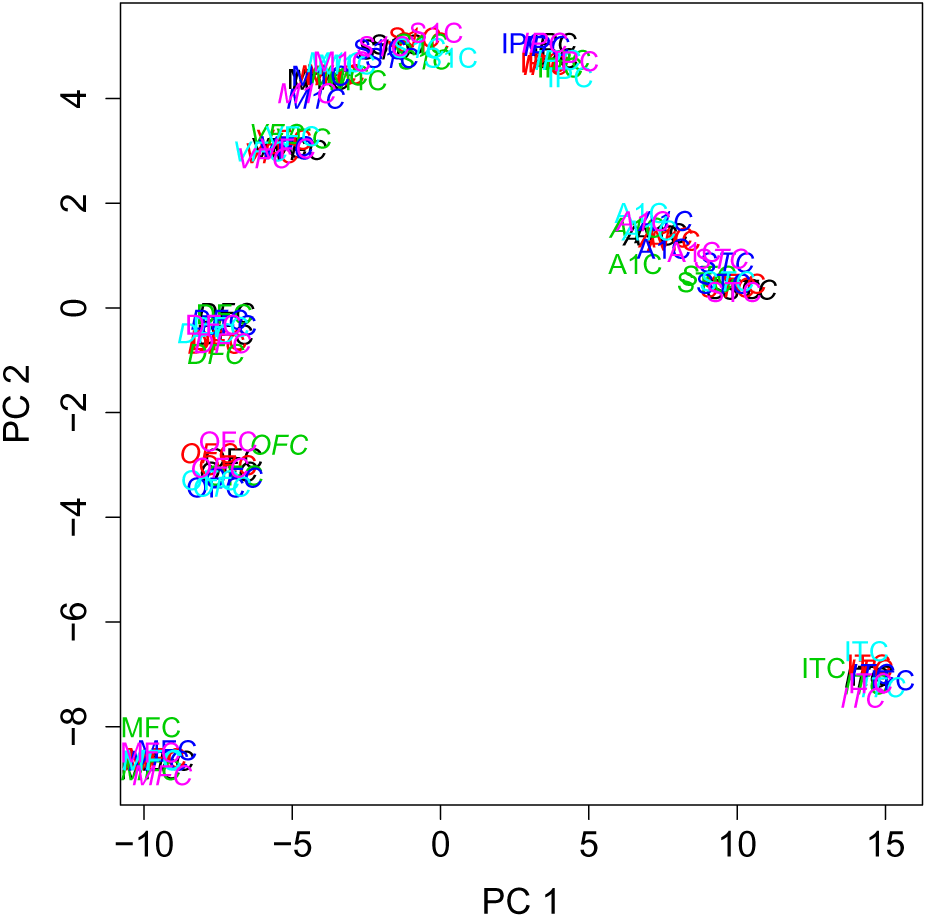
Window l, AC-PCA, λ = 20

**Table 1:** Span of the windows in this study.

| Window | Age |
| --- | --- |
| 1 | $12\text{PCW} \leq \text{Age} < 16\text{PCW}$ |
| 2 | $16\text{PCW} \leq \text{Age} < 21\text{PCW}$ |
| 3 | $21\text{PCW} \leq \text{Age} < 35\text{PCW}$ |
| 4 | $35\text{PCW} \leq \text{Age} < 10\text{M}$ |
| 5 | $10\text{M} \leq \text{Age} < 8\text{Y}$ |
| 6 | $8\text{Y} \leq \text{Age} < 19\text{Y}$ |
| 7 | $19\text{Y} \leq \text{Age} < 30\text{Y}$ |
| 8 | $30\text{Y} \leq \text{Age} < 42\text{Y}$ |
| 9 | $42\text{Y} \leq \text{Age}$ |

M: postnatal months
PCW: post-conception weeks
Y: postnatal years.

**Table 2:**
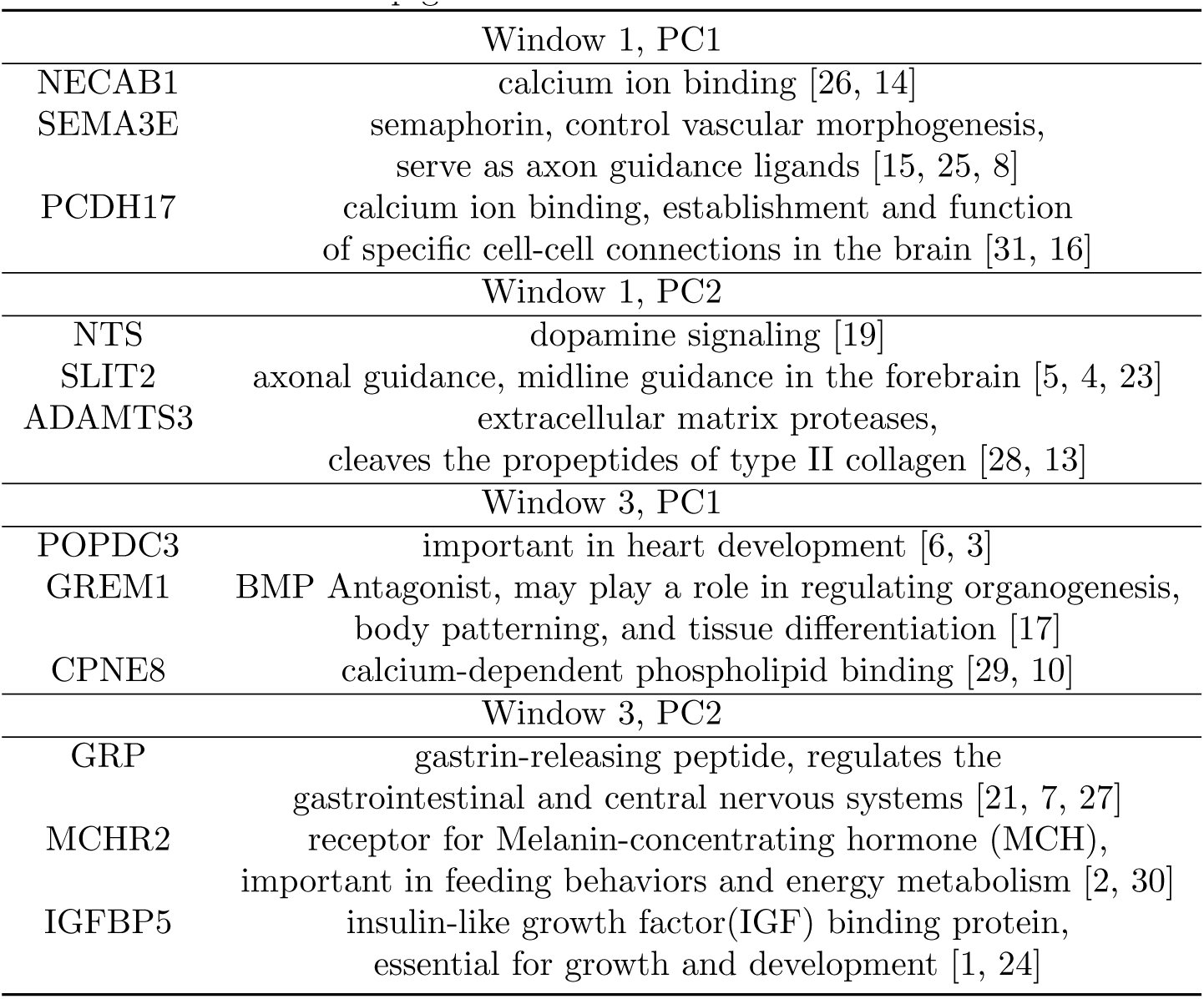
Top genes in the PCs and their functions.

## References

[1] Jess Allen, Hazel M Davey, David Broadhurst, Jim K Heald, Jem J Rowland, Stephen G Oliver, and Douglas B Kell. High-throughput classification of yeast mutants for functional genomics using metabolic footprinting. Nature biotechnology, 21(6):692–696, 2003.

[2] Elnaz Barshan, Ali Ghodsi, Zohreh Azimifar, and Mansoor Zolghadri Jahromi. Supervised principal component analysis: Visualization, classification and regression on subspaces and submanifolds. Pattern Recognition, 44(7):1357–1371, 2011.

[3] Susan E Celniker, Laura AL Dillon, Mark B Gerstein, Kristin C Gunsalus, Steven Henikoff, Gary H Karpen, Manolis Kellis, Eric C Lai, Jason D Lieb, David M MacAlpine, et al. Unlocking the secrets of the genome. Nature, 459(7249):927–930, 2009.

[4] Spyros Darmanis, Steven A Sloan, Ye Zhang, Martin Enge, Christine Caneda, Lawrence M Shuer, Melanie G Hayden Gephart, Ben A Barres, and Stephen R Quake. A survey of human brain transcriptome diversity at the single cell level. Proceedings of the National Academy of Sciences, page 201507125, 2015.

[5] Rajdeep Das, Nevenka Dimitrova, Zhenyu Xuan, Robert A Rollins, Fatemah Haghighi, John R Edwards, Jingyue Ju, Timothy H Bestor, and Michael Q Zhang. Computational prediction of methylation status in human genomic sequences. Proceedings of the National Academy of Sciences, 103(28):10713–10716, 2006.

[6] Johann A Gagnon-Bartsch and Terence P Speed. Using control genes to correct for unwanted variation in microarray data. Biostatistics, 13(3):539–552, 2012.

[7] Mark B Gerstein, Joel Rozowsky, Koon-Kiu Yan, Daifeng Wang, Chao Cheng, James B Brown, Carrie A Davis, LaDeana Hillier, Cristina Sisu, Jingyi Jessica Li, et al. Comparative analysis of the transcriptome across distant species. Nature, 512(7515):445–448, 2014.

[8] Thomas J Giordano, Rork Kuick, Tobias Else, Paul G Gauger, Michelle Vinco, Juliane Bauersfeld, Donita Sanders, Dafydd G Thomas, Gerard Doherty, and Gary Hammer. Molecular classification and prognostication of adrenocortical tumors by transcriptome profiling. Clinical Cancer Research, 15(2):665–676, 2009.

[9] Arthur Gretton, Olivier Bousquet, Alex Smola, and Bernhard Scholkopf. Measuring statistical dependence with hilbert-schmidt norms. In Algorithmic learning theory, pages 63–77. Springer, 2005.

[10] W Evan Johnson, Cheng Li, and Ariel Rabinovic. Adjusting batch effects in microarray expression data using empirical bayes methods. Biostatistics, 8(1):115–127, 2007.

[11] Iain M Johnstone and Arthur Yu Lu. On consistency and sparsity for principal components analysis in high dimensions. Journal of the American Statistical Association, 104(456), 2009.

[12] Ian Jolliffe. Principal component analysis. Wiley Online Library, 2002.

[13] Ian T Jolliffe, Nickolay T Trendafilov, and Mudassir Uddin. A modified principal component technique based on the lasso. Journal of computational and Graphical Statistics, 12(3):531–547, 2003.

[14] Sungkyu Jung, JS Marron, et al. Pca consistency in high dimension, low sample size context. The Annals of Statistics, 37(6B):4104–4130, 2009.

[15] Hyo Jung Kang, Yuka Imamura Kawasawa, Feng Cheng, Ying Zhu, Xuming Xu, Mingfeng Li, Andre MM Sousa, Mihovil Pletikos, Kyle A Meyer, Goran Sedmak, et al. Spatio-temporal transcriptome of the human brain. Nature, 478(7370):453–459, 2011.

[16] Joseph B Kruskal and Myron Wish. Multidimensional scaling, volume 11. Sage, 197S.

[17] Marta Kulis, Simon Heath, Marina Bibikova, Ana C Queiros, Alba Navarro, Guillem Clot, Alejandra Martinez-Trillos, Giancarlo Castellano, Isabelle Brun-Heath, Magda Pinyol, et al. Epigenomic analysis detects widespread gene-body dna hypomethylation in chronic lymphocytic leukemia. Nature genetics, 44(11):1236–1242, 2012.

[18] Jeffrey T Leek, W Evan Johnson, Hilary S Parker, Andrew E Jaffe, and John D Storey. The sva package for removing batch effects and other unwanted variation in high-throughput experiments. Bioinformatics, 28(6):882–883, 2012.

[19] Jeffrey T Leek and John D Storey. Capturing heterogeneity in gene expression studies by surrogate variable analysis. PLoS Genet, 3(9):1724–1735, 2007.

[20] Jingyi Jessica Li, Haiyan Huang, Peter J Bickel, and Steven E Brenner. Comparison of d. melanogaster and c. elegans developmental stages, tissues, and cells by modencode rna-seq data. Genome research, 24(7):1086–1101, 2014.

[21] Zhixiang Lin, Stephan J Sanders, Mingfeng Li, Nenad Sestan, Hongyu Zhao, et al. A markov random field-based approach to characterizing human brain development using spatial-temporal transcriptome data. The Annals of Applied Statistics, 9(1):429–451, 2015.

[22] Zongming Ma et al. Sparse principal component analysis and iterative thresholding. The Annals of Statistics, 41(2):772–801, 2013.

[23] Jeremy A Miller, Song-Lin Ding, Susan M Sunkin, Kimberly A Smith, Lydia Ng, Aaron Szafer, Amanda Ebbert, Zackery L Riley, Joshua J Royall, Kaylynn Aiona, et al. Transcriptional landscape of the prenatal human brain. Nature, 508(7495):199–206, 2014.

[24] Hilary S Parker, Hector Corrada Bravo, and Jeffrey T Leek. Removing batch effects for prediction problems with frozen surrogate variable analysis. PeerJ, 2:e561, 2014.

[25] Mihovil Pletikos, Andre MM Sousa, Goran Sedmak, Kyle A Meyer, Ying Zhu, Feng Cheng, Mingfeng Li, Yuka Imamura Kawasawa, and Nenad Sestan. Temporal specification and bilaterality of human neocortical topographic gene expression. Neuron, 81(2):321–332, 2014.

[26] Xin Qi, Ruiyan Luo, and Hongyu Zhao. Sparse principal component analysis by choice of norm. Journal of multivariate analysis, 114:127–160, 2013.

[27] Markus Ringner. What is principal component analysis? Nature biotechnology, 26(3):303–304, 2008.

[28] Davide Risso, John Ngai, Terence P Speed, and Sandrine Dudoit. Normalization of rna-seq data using factor analysis of control genes or samples. Nature biotechnology, 32(9):896–902, 2014.

[29] Alexei A Sharov, Yulan Piao, Ryo Matoba, Dawood B Dudekula, Yong Qian, Vincent VanBuren, Geppino Falco, Patrick R Martin, Carole A Stagg, Uwem C Bassey, et al. Transcriptome analysis of mouse stem cells and early embryos. PLoS Biol, 1(3):E74, 2003.

[30] Daniela M Witten, Robert Tibshirani, and Trevor Hastie. A penalized matrix decomposition, with applications to sparse principal components and canonical correlation analysis. Biostatistics, page kxp008, 2009.

[31] Can Yang, Lin Wang, Shuqin Zhang, and Hongyu Zhao. Accounting for non-genetic factors by low-rank representation and sparse regression for eqtl mapping. Bioinformatics, 29(8):1026–1034, 2013.

[32] Ren Zhang and Yan Lin. Deg 5.0, a database of essential genes in both prokaryotes and eukaryotes. Nucleic acids research, 37(suppl 1):D455–D458, 2009.

[33] Hui Zou, Trevor Hastie, and Robert Tibshirani. Sparse principal component analysis. Journal of computational and graphical statistics, 15(2):265–286, 2007.

## References

[1] Susanne V Allander, Catharina Larsson, Ewa Ehrenborg, Adisak Suwanichkul, Gunther Weber, Sheila L Morris, Svetlana Bajalica, Michael C Kiefer, Holger Luthman, and David R Powell. Characterization of the chromosomal gene and promoter for human insulin-like growth factor binding protein-5. Journal of Biological Chemistry, 269(14):10891–10898, 1994.

[2] Songzhu An, Gene Cutler, Jack Jiagang Zhao, Shu-Gui Huang, Hui Tian, Wanbo Li, Lingming Liang, Miki Rich, Amy Bakleh, Juan Du, et al. Identification and characterization of a melanin-concentrating hormone receptor. Proceedings of the National Academy of Sciences, 98(13):7576–7581, 2001.

[3] Birgit Andrée, Tina Hillemann, Gania Kessler-Icekson, Thomas Schmitt-John, Harald Jockusch, Hans-Henning Arnold, and Thomas Brand. Isolation and characterization of the novel popeye gene family expressed in skeletal muscle and heart. Developmental biology, 223(2):371–382, 2000.

[4] Kim T Nguyen Ba-Charvet, Katja Brose, Le Ma, Kuan H Wang, Valerie Marillat, Constantino Sotelo, Marc Tessier-Lavigne, and Alain Chedotal. Diversity and specificity of actions of slit2 proteolytic fragments in axon guidance. The Journal of Neuroscience, 21(12):4281–4289, 2001.

[5] Kim Tuyen Nguyen Ba-Charvet, Katja Brose, Valerie Marillat, Tom Kidd, Corey S Goodman, Marc Tessier-Lavigne, Constantino Sotelo, and Alain Chedotal. Slit2-mediated chemorepulsion and collapse of developing forebrain axons. Neuron, 22(3):463–473, 1999.

[6] Thomas Brand. The popeye domain-containing gene family. Cell biochemistry and biophysics, 43(1):95–103, 2005.

[7] PL Brubaker and DJ Drucker. Minireview: glucagon-like peptides regulate cell proliferation and apoptosis in the pancreas, gut, and central nervous system. Endocrinology, 145(6):2653–2659, 2004.

[8] Sophie Chauvet, Samia Cohen, Yutaka Yoshida, Lylia Fekrane, Jean Livet, Odile Gayet, Louis Segu, Marie-Christine Buhot, Thomas M Jessell, Christopher E Henderson, et al. Gating of sema3e/plexind1 signaling by neuropilin-1 switches axonal repulsion to attraction during brain development. Neuron, 56(5):807–822, 2007.

[9] 1000 Genomes Project Consortium et al. A map of human genome variation from population-scale sequencing. Nature, 467(7319):1061–1073, 2010.

[10] Carl E Creutz, Jose L Tomsig, Sandra L Snyder, Marie-Christine Gautier, Feriel Skouri, Janine Beisson, and Jean Cohen. The copines, a novel class of c2 domain-containing, calciumdependent, phospholipid-binding proteins conserved from paramecium to humans. Journal of Biological Chemistry, 273(3):1393–1402, 1998.

[11] Fiona Cunningham, M Ridwan Amode, Daniel Barrell, Kathryn Beal, Konstantinos Billis, Simon Brent, Denise Carvalho-Silva, Peter Clapham, Guy Coates, Stephen Fitzgerald, et al. Ensembl 2015. Nucleic acids research, 43(D1):D662–D669, 2015.

[12] John Duchi, Shai Shalev-Shwartz, Yoram Singer, and Tushar Chandra. Efficient projections onto the l 1-ball for learning in high dimensions. In Proceedings of the 25th international conference on Machine learning, pages 272–279. ACM, 2008.

[13] Russell J Fernandes, Satoshi Hirohata, J Michael Engle, Alain Colige, Daniel H Cohn, David R Eyre, and Suneel S Apte. Procollagen ii amino propeptide processing by adamts-3 insights on dermatosparaxis. Journal of Biological Chemistry, 276(34):31502–31509, 2001.

[14] Alexandra P Few, Nathan J Lautermilch, Ruth E Westenbroek, Todd Scheuer, and William A Catterall. Differential regulation of cav2. 1 channels by calcium-binding protein 1 and visinin-like protein-2 requires n-terminal myristoylation. The Journal of neuroscience, 25(30):7071–7080, 2005.

[15] Chenghua Gu, Yutaka Yoshida, Jean Livet, Dorothy V Reimert, Fanny Mann, Janna Merte, Christopher E Henderson, Thomas M Jessell, Alex L Kolodkin, and David D Ginty. Semaphorin 3e and plexin-d1 control vascular pattern independently of neuropilins. Science, 307(5707):265–268, 2005.

[16] Naosuke Hoshina, Asami Tanimura, Miwako Yamasaki, Takeshi Inoue, Ryoji Fukabori, Teiko Kuroda, Kazumasa Yokoyama, Tohru Tezuka, Hiroshi Sagara, Shinji Hirano, et al. Protocadherin 17 regulates presynaptic assembly in topographic corticobasal ganglia circuits. Neuron, 78(5):839–854, 2013.

[17] David R Hsu, Aris N Economides, Xiaorong Wang, Peter M Eimon, and Richard M Harland. The xenopus dorsalizing factor gremlin identifies a novel family of secreted proteins that antagonize bmp activities. Molecular cell, 1(5):673–683, 1998.

[18] W Evan Johnson, Cheng Li, and Ariel Rabinovic. Adjusting batch effects in microarray expression data using empirical bayes methods. Biostatistics, 8(1):118–127, 2007.

[19] Patrick Kitabgi. Neurotensin modulates dopamine neurotransmission at several levels along brain dopaminergic pathways. Neurochemistry international, 14(2):111–119, 1989.

[20] Jeffrey T Leek and John D Storey. Capturing heterogeneity in gene expression studies by surrogate variable analysis. PLoS Genet, 3(9):1724–1735, 2007.

[21] TJ McDonald, H Jornvall, G Nilsson, M Vagne, M Ghatei, SR Bloom, and V Mutt. Characterization of a gastrin releasing peptide from porcine non-antral gastric tissue. Biochemical and biophysical research communications, 90(1):227–233, 1979.

[22] Hilary S Parker, Hector Corrada Bravo, and Jeffrey T Leek. Removing batch effects for prediction problems with frozen surrogate variable analysis. PeerJ, 2:e561, 2014.

[23] Andrew S Plump, Lynda Erskine, Christelle Sabatier, Katja Brose, Charles J Epstein, Corey S Goodman, Carol A Mason, and Marc Tessier-Lavigne. Slitl and slit2 cooperate to prevent premature midline crossing of retinal axons in the mouse visual system. Neuron, 33(2):219–232, 2002.

[24] Dervis AM Salih, Gyanendra Tripathi, Cathy Holding, Tadge AM Szestak, M Ivelisse Gonzalez, Emma J Carter, Laura J Cobb, Joan E Eisemann, and Jennifer M Pell. Insulin-like growth factor-binding protein 5 (igfbp5) compromises survival, growth, muscle development, and fertility in mice. Proceedings of the National Academy of Sciences, 101(12):4314–4319, 2004.

[25] Guido Serini, Donatella Valdembri, Sara Zanivan, Giulia Morterra, Constanze Burkhardt, Francesca Caccavari, Luca Zammataro, Luca Primo, Luca Tamagnone, Malcolm Logan, et al. Class 3 semaphorins control vascular morphogenesis by inhibiting integrin function. Nature, 424(6947):391–396, 2003.

[26] S Sugita, A Ho, and TC Südhof. Necabs: A family of neuronal ca 2+- binding proteins with an unusual domain structure and a restricted expression pattern. Neuroscience, 112(1):51–63, 2002.

[27] Yan-Gang Sun and Zhou-Feng Chen. A gastrin-releasing peptide receptor mediates the itch sensation in the spinal cord. Nature, 448(7154):700–703, 2007.

[28] Bor Luen Tang. Adamts: a novel family of extracellular matrix proteases. The international journal of biochemistry & cell biology, 33(1):33–44, 2001.

[29] Jose Luis Tomsig and Carl E Creutz. Biochemical characterization of copine: a ubiquitous ca2+-dependent, phospholipid-binding protein. Biochemistry, 39(51):16163–16175, 2000.

[30] Suke Wang, Jiang Behan, Kim O’Neill, Blair Weig, Steven Fried, Thomas Laz, Marvin Bayne, Eric Gustafson, and Brian E Hawes. Identification and pharmacological characterization of a novel human melanin-concentrating hormone receptor, mch-r2. Journal of Biological Chemistry, 276(37):34664–34670, 2001.

[31] Takeshi Yagi and Masatoshi Takeichi. Cadherin superfamily genes: functions, genomic organization, and neurologic diversity. Genes & development, 14(10):1169–1180, 2000.

